# Modifiable risk factors moderate the relationship between beta-amyloid and cognition in midlife

**DOI:** 10.1101/229062

**Authors:** Lindsay R. Clark, Rebecca L. Koscik, Samantha L. Allison, Sara E. Berman, Cynthia M. Carlsson, Derek Norton, Barbara B. Bendlin, Sanjay Asthana, Tobey Betthauser, Bradley T. Christian, Sterling C. Johnson

## Abstract

Although evidence suggests a relationship between elevated beta-amyloid and cognitive decline, approximately 30% of older adults with positive markers of amyloid remain cognitively healthy. Our objective was to test if the presence of modifiable risk factors (i.e., central obesity, hypertension, and depressive symptoms) moderated the relationship between amyloid and longitudinal cognitive performance. Data were from 207 adults (140 females; age range=40-70) enriched for Alzheimer’s disease risk (73% parental history of Alzheimer’s disease) enrolled in the Wisconsin Registry for Alzheimer’s Prevention study. Participants completed at least three neuropsychological evaluations and one biomarker visit ([C11]Pittsburgh Compound B PET scan or lumbar puncture). Participants were characterized as high or low on beta-amyloid using cutoffs developed for [C11]Pittsburgh Compound B-PET distribution volume ratio or CSF amyloid beta 1-42 values. Participants were also coded as high or low risk on obesity (waist circumference > 102 cm for males or 88 cm for females), hypertension (systolic blood pressure ≥ 140 mm Hg or diastolic blood pressure ≥ 90 mm Hg), and depressive symptoms (Center for Epidemiologic Studies of Depression scale ≥ 16). Linear mixed effects regression models examined three-way interactions between modifiable risk factor status x beta-amyloid status x visit age on longitudinal Verbal Learning & Memory and Speed & Flexibility factor scores. Results indicated that the relationship between beta-amyloid and Verbal Learning & Memory decline was moderated by the presence of hypertension at baseline (*p* = .02), presence of hypertension at all visits (*p* = .001), and presence of obesity at all visits (*p* = .049). Depressive symptoms did not moderate the association between beta-amyloid and longitudinal Verbal Learning & Memory (*p* = .62) or Speed & Flexibility (*p* = .15) performances. In this at-risk for Alzheimer’s disease cohort, modifiable risk factors of hypertension and obesity moderated the relationship between beta-amyloid and cognitive decline. Identification and modification of these risk factors in late middle age may slow the effect of amyloid on the progression of cognitive symptoms.

## INTRODUCTION

Although available treatments for Alzheimer’s disease may provide some short-term benefits, they have limited efficacy in terms of modifying the course of the disease (Klafki *et al.*, 2006; Salomone *et al.*, 2012). The lack of an effective disease-modifying medication has led to increased efforts targeted at both early detection of Alzheimer’s disease and modifiable risk factors that may influence disease progression. Two commonly used ante-mortem biomarkers allowing for early detection of Alzheimer’s disease-related pathology are the PET imaging tracer [C11]Pittsburgh Compound B, which allows for imaging of amyloid deposition in vivo (Ikonomovic *et al.*, 2008), and CSF levels of amyloid-beta 1-42, which are correlated with the formation of amyloid plaques in the brain (Strozyk *et al.*, 2003; Fagan *et al.*, 2006). Prior investigations indicate that cognitively normal adults with higher mean cortical binding potential values for the PET imaging tracer [C11]Pittsburgh Compound B and/or lower cerebrospinal levels of amyloid beta 1-42 are at increased risk of developing dementia (Morris *et al.*, 2009; Roe *et al.*, 2011; Soldan *et al.*, 2013; Chen *et al.*, 2014). Furthermore, cognitively healthy adults with elevated beta-amyloid deposition are more likely to exhibit cognitive decline over time than adults with lower beta-amyloid levels (Gustafson *et al.*, 2007; Lim *et al.*, 2014; Ossenkoppele *et al.*, 2014; Clark *et al.*, 2016; Petersen *et al.*, 2016). Although elevated amyloid deposition is associated with increased risk of both cognitive decline and dementia, up to 30% of older adults with elevated amyloid deposition remain cognitively normal in late life (Morris *et al.*, 2010). This finding suggests that not all adults with elevated amyloid deposition will progress to dementia, and that other factors may moderate the relationship between amyloid deposition and cognitive decline.

Supporting this hypothesis, epidemiological studies suggest that seven potentially modifiable risk factors for Alzheimer’s disease, including midlife hypertension, midlife obesity, smoking, depression, low educational attainment, physical inactivity, and diabetes may account for up to half of dementia cases in the United States (Barnes and Yaffe, 2011). Several studies report greater risk for dementia and/or longitudinal decline on neuropsychological measures in adults with depressive symptomatology (Berger *et al.*, 1999; Green *et al.*, 2003; Sachs-Ericsson *et al.*, 2005; Ownby *et al.*, 2006; Royall and Palmer, 2013; Verdelho *et al.*, 2013; Geda *et al.*, 2014; Xu *et al.*, 2015), obesity (Kivipelto *et al.*, 2005; Whitmer *et al.*, 2005a; Wolf *et al.*, 2007; Sabia *et al.*, 2009; Profenno *et al.*, 2010; Dahl *et al.*, 2013; Xu *et al.*, 2015), or hypertension (Whitmer *et al.*, 2005b; Wolf *et al.*, 2007; Gao *et al.*, 2009; Gottesman *et al.*, 2014; Haring *et al.*, 2015; Walker *et al.*, 2017). Moreover, prior investigations revealed elevated amyloid deposition in non-demented older adults with depression (Harrington *et al.*, 2015) or hypertension (Langbaum *et al.*, 2012; Nation *et al.*, 2013; Hughes *et al.*, 2014). Two recent studies also observed a relationship between midlife obesity and amyloid deposition in later life (Chuang *et al.*, 2016; Gottesman *et al.*, 2017). Finally, one prior investigation demonstrated that adults with abnormal plasma amyloid levels and elevated blood pressure at midlife were at greatest risk of developing Alzheimer’s disease (Shah *et al.*, 2012); however, no study to date has examined whether cognitively healthy middle-aged adults with these risk factors and elevated amyloid deposition are more likely to decline than those with elevated amyloid but absence of these risk factors.

Therefore, the purpose of the current study was to determine if the presence of modifiable risk factors moderate the relationship between amyloid deposition and longitudinal performance on neuropsychological measures in cognitively normal late middle-aged adults. We decided to focus this study on three risk factors that were objectively measured and well-characterized in the Wisconsin Registry for Alzheimer’s Prevention cohort: hypertension, obesity, and depression. We investigated the moderating effects of these risk factors on the relationship between amyloid deposition (as assessed via the PET imaging tracer [C11]Pittsburgh Compound B or cerebrospinal levels of amyloid beta 1-42) and longitudinal neuropsychological performance. We hypothesized that each risk factor would moderate the relationship between amyloid burden and longitudinal cognitive performance (e.g., three-way interaction among visit age (time-varying) x risk factor group (time-invariant) x amyloid group (time-invariant) would account for a significant amount of variance in longitudinal neuropsychological performance).

## MATERIALS AND METHODS

### Participants

Data were from 207 participants enrolled in the Wisconsin Registry for Alzheimer’s Prevention (WRAP) study, which consists of a cohort of ~1550 asymptomatic (at study entry) late middle-aged adults enriched for parental history of Alzheimer’s disease (Johnson *et al.*, 2018). The ongoing parent study includes biennial evaluations that involve a physical exam, labs, a neuropsychological evaluation, and optional linked studies for acquisition of neuroimaging and CSF biomarkers of Alzheimer’s disease. The Wisconsin Registry for Alzheimer’s Prevention protocol includes a baseline neuropsychological evaluation, a second visit four years after baseline, and subsequent visits every two years. Inclusion criteria for this study were as follows: outcome data for at least three study visits, no diagnosis of dementia, and completion of either an amyloid PET scan or lumbar puncture. The inclusion of human subjects in this study was approved by the University of Wisconsin-Madison Institutional Review Board and all participants provided informed consent.

### Modifiable Risk Factor Assessment

The modifiable risk factors included in this study were hypertension, obesity, and depression, which were chosen because they were included in the epidemiological study previously described (Barnes and Yaffe, 2011) and were present in >10% of the entire WRAP cohort. Blood pressure and anthropometric measures were obtained at study visit 2 and subsequent visits according to the Atherosclerosis Risk in Communities Study protocol. Prior to cognitive test administration, participants were instructed to sit for 10 minutes and then have blood pressure readings obtained. Blood pressure was measured up to three times within an examination visit to obtain a stable measure, with the participant seated using a random-zero sphygmomanometer. Cuff size was chosen appropriate to the participant’s arm circumference. Hypertension was defined according to the guidelines of the Joint National Committee on Detection, Evaluation, and Treatment of High Blood Pressure (i.e., systolic blood pressure ≥ 140 mm Hg or diastolic blood pressure ≥ 90 mm Hg (James *et al.*, 2014)). Use of antihypertensive medication was included as a covariate in primary analyses and examined further in post-hoc analyses. Waist circumference was measured with an anthropometric tape to the nearest centimeter with the participant standing. The waist circumference was taken at the level of the natural waist (narrowest part). Waist measurements were taken twice by trained staff and the smallest measurement was used. Waist circumference measurements greater than 88 cm for women and 102 cm for men were coded as obese based on standard guidelines (WHO, 2011). Depressive symptoms were assessed using the Center for Epidemiologic Studies of Depression scale, with total scores ≥ 16 coded as depressed (Lewinsohn *et al.*, 1997).

### Cognitive Assessment

A comprehensive neuropsychological assessment was completed at each visit (Johnson *et al.*, 2018). A prior factor analysis indicated that learning trials 3-5 and the delayed recall trial (trial 7) on the Rey Auditory Verbal Learning Test loaded onto a Verbal Learning & Memory factor and that measures of speed and executive function (Trailmaking Test Parts A & B, Stroop Color-Word Interference condition) loaded onto a Speed & Flexibility factor (Dowling *et al.*, 2010). Each factor was calculated as a weighted composite of the contributing tests and then standardized around the baseline mean and standard deviation of the composite (i.e., each factor is a z-score) (Koscik *et al.*, 2014). These factors were selected for analyses because they represent cognitive domains that have been shown to exhibit early decline and associations with beta-amyloid in preclinical Alzheimer’s disease in prior meta-analyses (Backman *et al.*, 2005; Hedden *et al.*, 2013; Duke Han *et al.*, 2017), and because they include measures that were given at all study visits.

### Amyloid status determination

Positive or negative amyloid status was determined by mean distribution volume ratio obtained from the PET imaging tracer [C11]Pittsburgh Compound B on a Siemens EXACT HR+ scanner or by amyloid beta 1-42 or amyloid beta 1-42/amyloid beta 1-40 levels in CSF obtained from a lumbar puncture.

Detailed methods for [C-11]PiB radiochemical synthesis, [C11]Pittsburgh Compound B-PET sequence parameters, and distribution volume ratio map generation have been described previously (Johnson *et al.*, 2014). Eight bilateral regions of interest (angular gyrus, anterior cingulate gyrus, posterior cingulate gyrus, frontal medial orbital gyrus, precuneus, supramarginal gyrus, middle temporal gyrus, and superior temporal gyrus) were selected from the automated anatomic labeling atlas, standardized, and reverse warped to native space. The mean distribution volume ratio across these eight regions of interest was calculated. Similar to prior studies in this cohort, a mean distribution volume ratio of 1.19 was used to define PET amyloid positivity (Racine *et al.*, 2016).

Cerebrospinal fluid was collected in the morning after a minimum 12-hour fast. A Sprotte spinal needle was used to extract twenty-two mL of cerebrospinal fluid from the L3-L4 or L4-L5 vertebral interspace via gentle extraction into polypropylene syringes. Within 30 minutes of collection, the cerebrospinal fluid was combined, gently mixed, centrifuged to remove red blood cells or other debris, aliquoted into 0.5-mL polypropylene tubes, and stored at −80°C. Samples were analyzed at the Clinical Neurochemistry Laboratory at the Sahlgrenska Academy of the University of Gothenburg, Sweden for amyloid beta 1-40 and amyloid beta 1-42 using commercially available enzyme-linked immunosorbent assay methods (INNOTEST assays, Fujirebio, Ghent, Belgium; Triplex assays, MSD Human Aβ peptide ultra-sensitive kit, Meso Scale Discovery, Gaithersburg, MD). An amyloid beta 1-42 (innotest) value < 471.54 or an amyloid beta 1-42/amyloid beta 1-40 < .09 was used to define amyloid positivity, based on prior receiver operating characteristic analyses showing these values best discriminated cognitively healthy adults from individuals with dementia (Clark *et al.*, Under Review).

### Statistical Analysis

Blinding and randomization were not performed for this study as this was a retrospective analysis of an observational cohort. Chi-square analyses were conducted in SPSS version 24 to test the relationships between the three modifiable risk factors, and the relationships between each modifiable risk factor and amyloid status. To test the hypothesis that longitudinal cognitive performance would vary by risk factor and amyloid status, linear mixed effects models were conducted in R version 3.3.1 using the lme4 package version 1.1-12. Outcome variables were Verbal Learning & Memory and Speed & Flexibility factor scores. Random effects included intercept and slope nested within-subject. Fixed effects for model 1 included hypertension status, amyloid status, age at each visit (time-varying variable; centered on the sample mean), hypertension status x amyloid status, age at each visit x hypertension status, age at each visit x amyloid status, and age at each visit x hypertension status x amyloid status. Fixed effects for models 2 and 3 were identical to model 1 with the exception of obesity (yes/no) and depression (yes/no) status instead of hypertension. Covariates included age at biomarker visit, sex, education, practice effects (number of prior exposures to cognitive test [total visits completed −1]), and treatment status (e.g., antihypertensive medication use (yes vs. no) for model 1 or antidepressant medication use (yes vs. no) for model 3). The overall significance of the three-way interaction term was assessed by likelihood ratio tests comparing the full model and a nested model that did not include the three-way interaction term. Statistical comparison of model coefficients to determine direction of group differences was performed using the Wald test. Statistical tests were two-tailed (except likelihood ratio test which was one-tailed) and an alpha-level of *p* < .05 was used to determine statistical significance. Model fit was evaluated by visual inspection of the residuals and a Pearson goodness-of-fit test. In primary models, each risk factor variable was based on status at study visit 2 (blood pressure and waist circumference were not acquired using the standard protocol at visit 1). Secondary models included cumulative risk factor information (and medication information) from all visits (e.g., yes = risk factor present at any visit; no = risk factor absent at all visits). For hypertension and depression, we also examined whether cognitive trajectories varied across groups defined by combining objective measurement of the risk factor and medication use (4 levels: non-symptomatic/non-treated, non-symptomatic/treated, symptomatic/non-treated, symptomatic/treated). Models run in primary analyses were re-run replacing the original visit 2 binary risk factor and medication covariate with this 4-level risk factor.

## RESULTS

### Sample characteristics

The sample was on average 55 years of age at baseline, 59 years of age at visit 2 (risk factor data acquisition), and 61 years of age at biomarker visit. The sample was 68% female, highly educated (mean 16 years of education), and enriched for Alzheimer’s disease risk (73% had at least one parent with Alzheimer’s disease; 38% apolipoprotein (*APOE*) ε4 genotype carriers; see Table 1). All 207 participants completed at least three neuropsychological evaluations. Twenty-five participants (12%) completed three study visits (six years follow-up), 82 (40%) completed four study visits (eight years follow-up), and 100 (48%) completed five study visits (ten years follow-up). Sixty-two participants (30%) were beta-amyloid positive. At visit 2, *n* = 35 (17%) were hypertensive, *n* = 81 (39%) were obese, and *n* = 14 (7%) were depressed. Across all visits available, *n* = 77 (37%) were hypertensive at one or more visits, *n* = 112 (54%) were obese at one or more visits, and *n* = 34 (16%) were depressed at one or more visits. Results of chi-square analyses indicated a significant association between hypertension and obesity status at visit 2 (*χ*^*2*^*(1)* = 7.70, *p* < .01) with 21 of the 35 hypertensive participants also meeting criteria for obesity. Similar results were observed when risk factor status was defined across all visits (*χ*^*2*^*(1)* = 7.26, *p* < .01), with 51 of the 77 hypertensive participants also meeting criteria for obesity. There was no significant relationship between hypertension and depression status at visit 2 (*χ*^*2*^*(1)* = 3.06, *p* = .08), with all 14 of the depressed participants in the non-hypertensive group. Lastly, there was no significant relationship between obesity and depression at visit 2 (*χ*^*2*^*(1)* = 0.09, *p* = .77), with about half of the depressed group (*n* = 6, 43%) also meeting criteria for obesity. Similar results were observed when risk factor status was defined based on data across all visits available.

**Table 1.** Sample characteristics

| Characteristics | Mean (SD) or N (%) |
| --- | --- |
| Baseline age | 54.6 (6.3) |
| Biomarker (PET or CSF) age | 60.7 (6.4) |
| Sex (Female) | 140 (68%) |
| Education (years) | 16.1 (2.4) |
| *APOE* (ε4 carrier) | 78 (38%) |
| Parental history of Alzheimer's disease (positive) | 151 (73%) |
| Baseline Verbal Learning & Memory (z-score) | .08 (1.0) |
| Baseline Speed & Flexibility (z-score) | -0.003 (0.9) |
| Beta-amyloid positive | 62 (30%) |
| Beta-amyloid modality acquired | CSF: 38 (18%) PiB: 84 (41%)<br>Both CSF and PiB: 85 (41%) |
| Systolic blood pressure (wave 2) | 124.5 (15.3) |
| Diastolic blood pressure (wave 2) | 74.4 (9.5) |
| Waist circumference (wave 2) | 91.6 (15.4) |
| Center for Epidemiologic Studies of Depression scale<br>total score (wave 2) | 6.2 (5.9) |
| Anti-hypertensive medication use (wave 2) | 40 (19%) |
| Anti-hypertensive medication use at least one visit<br>(cumulative) | 66 (32%) |

### Relationship between modifiable risk factors and beta-amyloid status

Results of chi-square analyses indicated that risk factor status did not differ by amyloid status (see Table 2). Specifically, there were no differences in proportion of participants classified as amyloid positive by hypertension status at visit 2 (*χ*^*2*^*(1)* = .04, *p* = .83) or across all visits (*χ*^*2*^*(1)* = .37, *p* = .54), obesity status at visit 2 (*χ*^*2*^*(1)* = 1.03, *p* = .31) or across all visits (*χ*^*2*^*(1)* = 1.92, *p* = .17), or depression status at visit 2 (*χ*^*2*^*(1)* = .24, *p* = .63) or across all visits (*χ*^*2*^*(1)* = .55, *p* = .46).

**Table 2.**
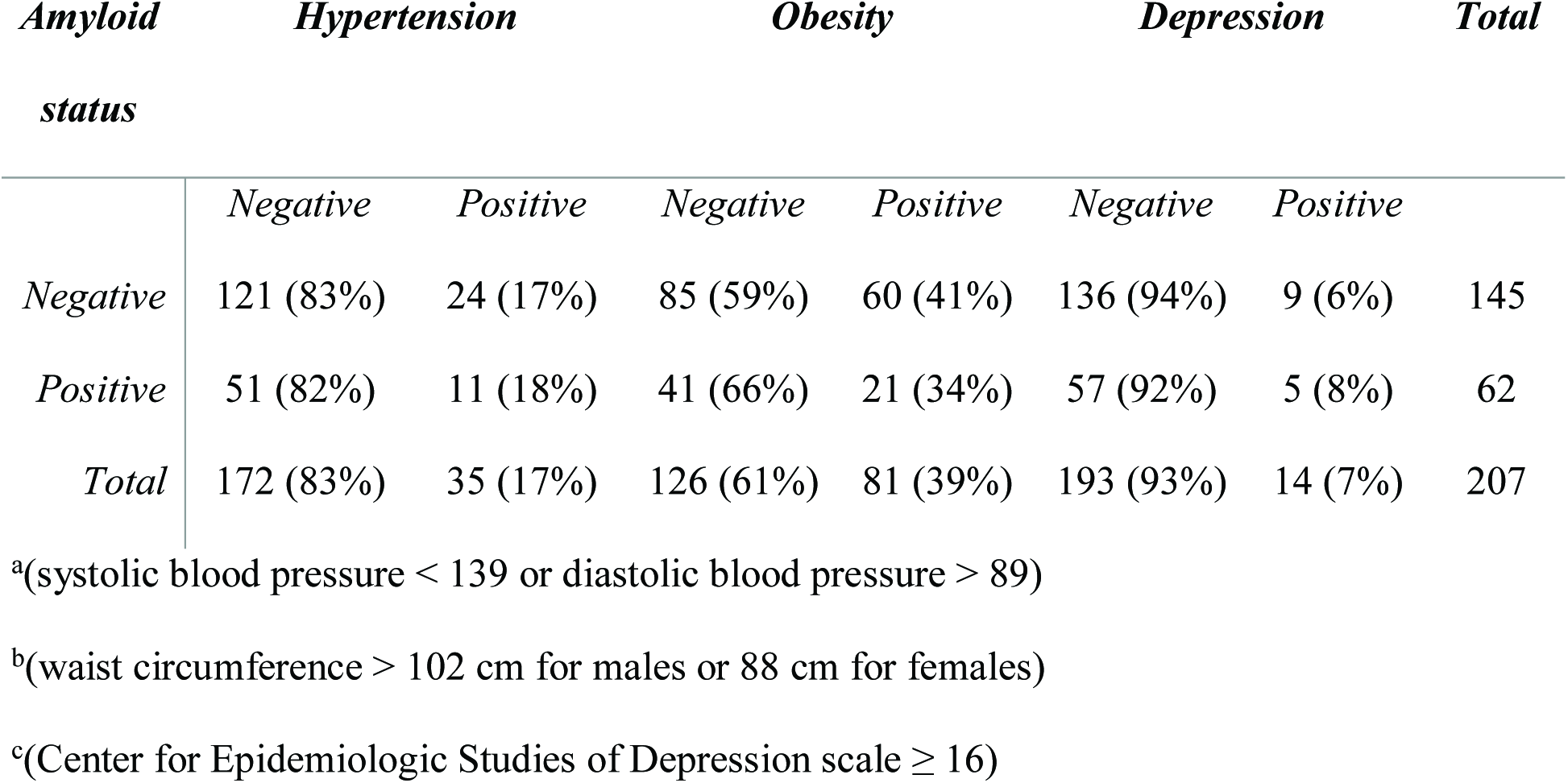
Proportion of sample classified as amyloid negative and positive within each risk factor group

| *Amyloid status* | *Hypertension* |  | *Obesity* |  | *Depression* |  | *Total* |
| --- | --- | --- | --- | --- | --- | --- | --- |
|  | *Negative* | *Positive* | *Negative* | *Positive* | *Negative* | *Positive* |  |
| *Negative* | 121 (83%) | 24 (17%) | 85 (59%) | 60 (41%) | 136 (94%) | 9 (6%) | 145 |
| *Positive* | 51 (82%) | 11 (18%) | 41 (66%) | 21 (34%) | 57 (92%) | 5 (8%) | 62 |
| *Total* | 172 (83%) | 35 (17%) | 126 (61%) | 81 (39%) | 193 (93%) | 14 (7%) | 207 |

<sup>a</sup>(systolic blood pressure < 139 or diastolic blood pressure > 89)
<sup>b</sup>(waist circumference > 102 cm for males or 88 cm for females)
<sup>c</sup>(Center for Epidemiologic Studies of Depression scale $\geq 16$ )

### Relationships among modifiable risk factors, beta-amyloid, and cognition

Regression diagnostics were performed and indicated that all models met the necessary assumptions. Specifically, model residuals appeared normally distributed, did not exhibit heteroscedasticity, and Pearson goodness-of-fit tests were non-significant. Random effects (e.g., intercept and slope) were not correlated with residuals and the random effect residuals were normally distributed.

#### Hypertension

Likelihood ratio tests comparing full and nested linear mixed-effects models indicated that the three-way interaction of hypertension status x amyloid status x visit age accounted for a statistically significant amount of variance in Verbal Learning & Memory performance (*χ*^*2*^*(1)* = 4.28, *p* = .04; see Table 3). This result indicates that the relationship between amyloid and rate of age-related decline in list-learning was also associated with hypertension status (see Figure 1 [Top]). Statistical comparison of model coefficients using Wald test indicated that those with elevated amyloid and hypertension (green line) did not exhibit significantly greater decline than those with elevated amyloid without hypertension (blue line) (*β* = −0.03 (SE=.03), *p* = .28), suggesting the significant interaction was driven instead by differences in decline between those with hypertension and elevated amyloid (green line) and those with hypertension and non-elevated amyloid (orange line) (*β* = −0.07 (SE=.03), *p* = .02). The interaction of hypertension status x amyloid status x visit age did not account for a significant amount of variability in Speed & Flexibility performance (*χ*^*2*^*(1)* = .09, *p* = .77; see Table 4).

**Figure 1 Legend.**
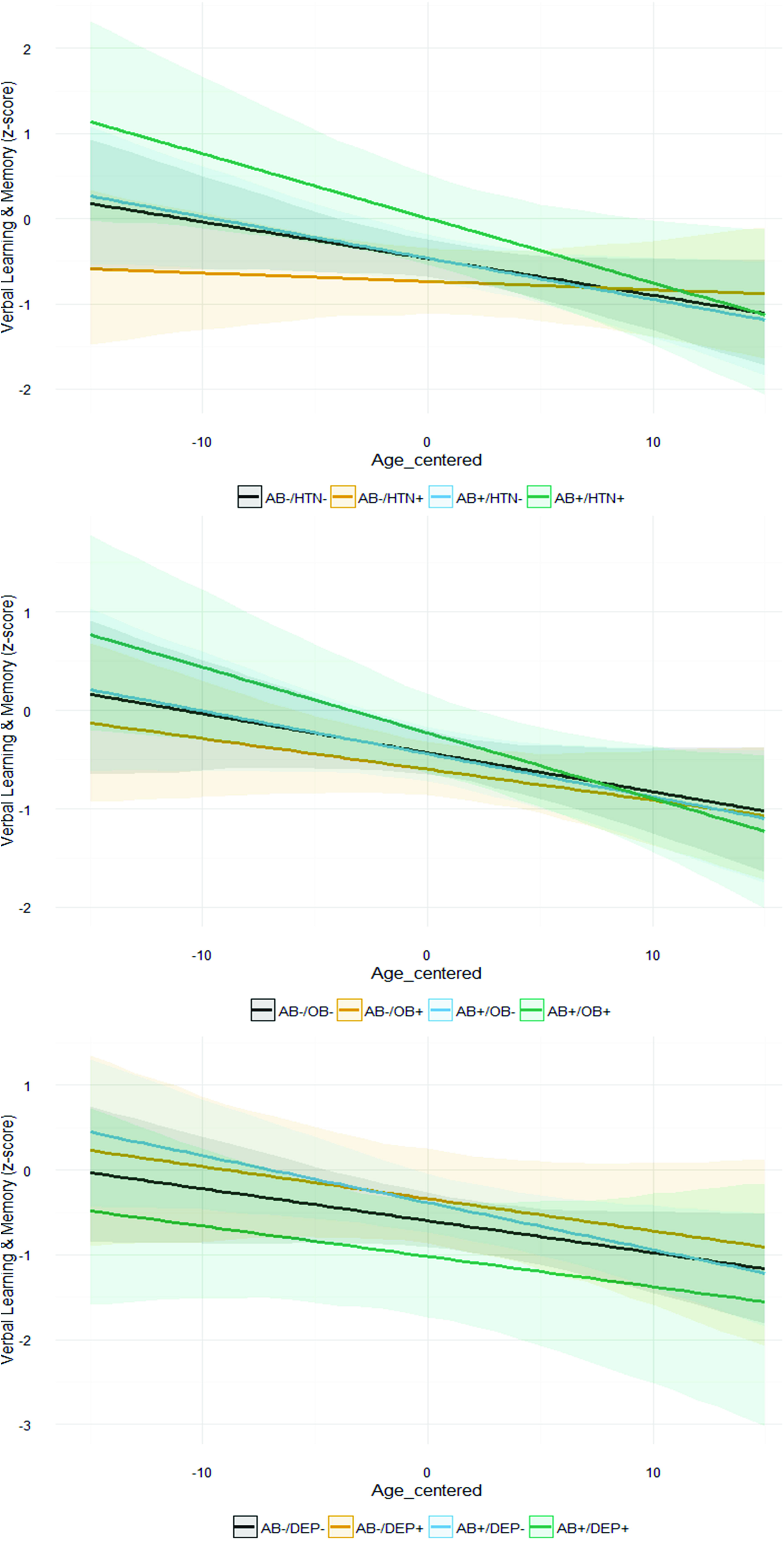
Graphs depict Verbal Learning & Memory z-scores on the y-axis, age at each visit (centered on mean age) on the x-axis, and estimated slopes for four beta-amyloid/risk factor groups adjusted for covariates of age at biomarker visit, sex, education, and practice effects. Risk factor groups are determined based on status at study visit 2. The top figure depicts the estimated slope for beta-amyloid negative and hypertension negative (black; *n* =121), beta-amyloid negative and hypertension positive (orange; *n* =24), beta-amyloid positive and hypertension negative (blue; *n* =51), and beta-amyloid positive and hypertension positive (green; *n* =11). The middle figure depicts the estimated slope for beta-amyloid negative and obesity negative (black; *n* =85), beta-amyloid negative and obesity positive (orange; *n* =60), beta-amyloid positive and obesity negative (blue; *n* =41), and beta-amyloid positive and obesity positive (green; *n* =21). The lower figure depicts the estimated slope for beta-amyloid negative and depression negative (black; *n* =136), beta-amyloid negative and depression positive (orange; *n* =9), beta-amyloid positive and depression negative (blue; *n* =57), and beta-amyloid positive and depression positive (green; *n* =5).

**Table 3.** Model parameter estimates for association between presence of modifiable risk factors at visit 2 and Verbal Learning & Memory outcome

|  | Risk Factor |  |  |
| --- | --- | --- | --- |
|  | Hypertension | Obesity | Depression |
| | $\beta$ (SE); [95% CI] | $\beta$ (SE); [95% CI] | $\beta$ (SE); [95% CI] |
| Intercept | -2.4 (1.6); [-5.5, 0.7] | -2.1 (1.6); [-5.1, 1.0] | -2.1 (1.6); [-5.1, 1.0] |
| Risk factor group | -0.3 (0.2); [-0.6, 0.1] | -0.2 (0.1); [-0.4, 0.1] | 0.3 (0.3); [-0.3, 0.8] |
| Amyloid group | 0.01 (0.1); [-0.3, 0.3] | -0.01 (0.1); [-0.3, 0.3] | 0.2 (0.1); [-0.04, 0.5] |
| Centered visit age (slope) | -0.04 (0.0); [-0.1, 0.0] | -0.04 (0.0); [-0.1, 0.01] | -0.04 (0.0); [-0.1, 0.01] |
| Biomarker age | 0.01 (0.0); [-0.04, 0.1] | 0.01 (0.0); [-0.04, 0.1] | -0.01 (0.0); [-0.04, 0.1] |
| Sex | **0.8\*\*\* (0.1); [0.6, 1.1]** | **0.8\*\*\* (0.1); [0.6, 1.1]** | **0.8\*\*\* (0.1); [0.6, 1.1]** |
| Education | **0.1\*\* (0.0); [0.02, 0.1]** | **0.1\*\* (0.0); [0.02, 0.1]** | **0.1\*\* (0.0); [0.02, 0.1]** |
| Practice effect | 0.1 (0.1); [-0.03, 0.2] | 0.1 (0.1); [-0.03, 0.2] | 0.1 (0.1); [-0.02, 0.2] |
| Treatment status | 0.1 (0.1); [-0.2, 0.3] | -- | -0.1 (0.1); [-0.4, 0.1] |
| Risk factor group x Amyloid group | **0.7\* (0.3); [0.1, 1.4]** | 0.4 (0.2); [-0.1, 0.9] | **-0.9\* (0.5); [-1.8, -0.003]** |
| Centered visit age x Risk factor group | **0.03\* (0.0); [0.0, 0.1]** | 0.01 (0.0); [-0.01, 0.03] | -0.0003 (0.0); [-0.1, 0.1] |
| Centered visit age x Amyloid group | -0.01 (0.0); [-0.03, 0.02] | -0.004 (0.0); [-0.03, 0.02] | -0.02 (0.0); [-0.04, 0.004] |
| Centered visit age x Amyloid group x Risk factor group | **-0.1\* (0.0); [-0.1, -0.002]** | -0.03 (0.0); [-0.1, 0.01] | 0.02 (0.0); [-0.1, 0.1] |

\*\*\*p ≤ .001; \*\*p ≤ .01; \*p ≤ .05; p-values for fixed effect coefficients were calculated using asymptomatic properties of the estimates

**Table 4.** Model parameter estimates for association between presence of modifiable risk factors at visit 2 and Speed & Flexibility outcome

|  | Risk Factor |  |  |
| --- | --- | --- | --- |
|  | Hypertension | Obesity | Depression |
| | $\beta$ (SE); [95% CI] | $\beta$ (SE); [95% CI] | $\beta$ (SE); [95% CI] |
| Intercept | 4.4** (1.4); [1.7, 7.1] | 4.3** (1.4); [1.6, 7.0] | 4.1** (1.4); [1.4, 6.8] |
| Risk factor group | 0.1 (0.2); [-0.2, 0.5] | 0.02 (0.1); [-0.2, 0.3] | -0.3 (0.3); [-0.8, 0.2] |
| Amyloid group | -0.1 (0.1); [-0.3, 0.2] | -0.003 (0.1); [-0.3, 0.3] | -0.1 (0.1); [-0.3, 0.2] |
| Age at each visit (centered) | 0.01 (0.0); [-0.03, 0.04] | 0.003 (0.0); [-0.04, 0.04] | -0.001 (0.0); [-0.04, 0.04] |
| Biomarker age | **-0.1\*\*\* (0.0); [-0.1, -0.03]** | **-0.1\*\*\* (0.0); [-0.1, -0.03]** | **-0.1\*\*\* (0.0); [-0.1, -0.03]** |
| Sex | -0.02 (0.1); [-0.3, 0.2] | -0.02 (0.1); [-0.2, 0.2] | -0.03 (0.1); [-0.2, 0.2] |
| Education | -0.001 (0.0); [-0.04, 0.04] | -0.001 (0.0); [-0.04, 0.04] | 0.0004 (0.0); [-0.04, 0.04] |
| Practice effect | 0.04 (0.1); [-0.1, 0.2] | 0.04 (0.1); [-0.1, 0.1] | 0.1 (0.1); [-0.1, 0.2] |
| Medication use | -0.04 (0.1); [-0.3, 0.2] | -- | 0.04 (0.1); [-0.2, 0.3] |
| Risk factor group x Amyloid group | -0.2 (0.3); [-0.7, 0.4] | -0.2 (0.2); [-0.6, 0.3] | -0.3 (0.4); [-1.1, 0.5] |
| Centered visit age x Risk factor group | -0.02 (0.0); [-0.04, 0.01] | -0.0004 (0.0); [-0.02, 0.02] | -0.01 (0.0); [-0.1, 0.04] |
| Centered visit age x Amyloid group | -0.01 (0.0); [-0.03, 0.004] | -0.01 (0.0); [-0.03, 0.01] | -0.01 (0.0); [-0.03, 0.01] |
| Centered visit age x Amyloid group x Risk factor group | 0.01 (0.0); [-0.04, 0.1] | -0.02 (0.0); [-0.1, 0.02] | -0.1 (0.0); [-0.1, 0.02] |

\*\*\* $p \leq .001$ ; \*\* $p \leq .01$ ; \* $p < .05$ ; p-values for fixed effect coefficients were calculated using asymptomatic properties of the estimates

Secondary analyses which replaced visit 2 hypertension status with cumulative hypertension status (e.g., yes if hypertensive at any visit (*n* = 77) and no if non-hypertensive at all visits), revealed similar results. Specifically, the three-way interaction of hypertension status x amyloid status x visit age accounted for a statistically significant amount of variance in Verbal Learning & Memory performance (*χ*^*2*^*(1)* = 10.29, *p* = .001; see Table 5), but not in Speed & Flexibility performance (*χ*^*2*^*(1)* = .18, *p* = .67; see Table 6). Statistical comparison of model coefficients indicated that those with amyloid positivity and hypertension (Figure 2 [Top green line]) exhibited significantly greater decline than those with amyloid positivity without hypertension (Figure 2 [Top blue line]) (*β* = −0.05 (SE=.02), *p* = .003).

**Table 5.** Model parameter estimates for association between presence of modifiable risk factors at any visit and Verbal Learning & Memory outcome

|  | Risk Factor |  |  |
| --- | --- | --- | --- |
|  | Hypertension | Obesity | Depression |
| | $\beta$ (SE); [95% CI] | $\beta$ (SE); [95% CI] | $\beta$ (SE); [95% CI] |
| Intercept | -2.0 (1.6); [-5.1, 1.0] | -2.1 (1.6); [-5.1, 1.0] | -2.2 (1.5); [-5.2, 0.8] |
| Risk factor group | -0.1 (0.1); [-0.3, 0.2] | -0.1 (0.1); [-0.3, 0.2] | 0.3 (0.2); [-0.1, 0.7] |
| Amyloid group | -0.03 (0.2); [-0.3, 0.3] | 0.1 (0.2); [-0.2, 0.4] | **0.3\* (0.1); [0.04, 0.6]** |
| Age at each visit (centered) | -0.04 (0.0); [-0.1, 0.004] | -0.04 (0.0); [-0.1, 0.01] | -0.04 (0.0); [-0.1, 0.01] |
| Biomarker age | 0.003 (0.0); [-0.04, 0.1] | 0.004 (0.0); [-0.04, 0.1] | 0.01 (0.0); [-0.04, 0.1] |
| Sex | **0.8\*\*\* (0.1); [0.6, 1.1]** | **0.8\*\*\* (0.1); [0.6, 1.1]** | **0.8\*\*\* (0.1); [0.6, 1.1]** |
| Education | **0.1\*\* (0.0); [0.02, 0.1]** | **0.1\*\* (0.0); [0.02, 0.1]** | **0.1\*\* (0.0); [0.02, 0.1]** |
| Practice effect | 0.1 (0.1); [-0.04, 0.2] | 0.1 (0.1); [-0.04, 0.2] | 0.1 (0.1); [-0.02, 0.2] |
| Medication use at some visits | 0.01 (0.1); [-0.3, 0.3] | -- | -0.2 (0.1); [-0.4, 0.1] |
| Medication use at all visits | 0.1 (0.2); [-0.3, 0.4] | -- | -0.1 (0.2); [-0.4, 0.2] |
| Risk factor group x Amyloid group | 0.4 (0.2); [-0.1, 0.9] | 0.03 (0.2); [-0.4, 0.5] | **-0.9\*\* (0.3); [-1.5, -0.3]** |
| Centered visit age x Risk factor group | 0.02 (0.0); [-0.01, 0.04] | 0.01 (0.0); [-0.01, 0.03] | -0.03 (0.0); [-0.1, 0.01] |
| Centered visit age x Amyloid group | 0.01 (0.0); [-0.01, 0.04] | 0.01 (0.0); [-0.02, 0.04] | -0.02 (0.0); [-0.04, 0.002] |
| Centered visit age x Amyloid group x Risk factor group | **-0.1\*\* (0.0); [-0.1, -0.03]** | **-0.04\* (0.0); [-0.1, .0002]** | 0.03 (0.0); [-0.02, 0.1] |

\*\*\* $p \leq .001$ ; \*\* $p \leq .01$ ; \* $p < .05$ ; p-values for fixed effect coefficients were calculated using asymptomatic properties of the estimates

**Table 6.** Model parameter estimates for association between presence of modifiable risk factors at any visit and Speed & Flexibility outcome

|  | Risk Factor |  |  |
| --- | --- | --- | --- |
|  | Hypertension | Obesity | Depression |
| | $\beta$ (SE); [95% CI] | $\beta$ (SE); [95% CI] | $\beta$ (SE); [95% CI] |
| Intercept | 4.5*** (1.4); [1.7, 7.2] | 4.4*** (1.4); [1.7, 7.0] | 4.2** (1.4); [1.5, 6.9] |
| Risk factor group | 0.1 (0.1); [-0.2, 0.4] | 0.02 (0.1); [-0.2, 0.3] | -0.1 (0.2); [-0.4, 0.3] |
| Amyloid group | -0.2 (0.1); [-0.4, 0.1] | 0.03 (0.2); [-0.3, 0.3] | -0.03 (0.1); [-0.3, 0.2] |
| Age at each visit (centered) | 0.01 (0.0); [-0.03, 0.04] | 0.01 (0.0); [-0.03, 0.1] | 0.002 (0.0); [-0.04, .04] |
| Biomarker age | **-0.1\*\*\* (0.0); [-0.1, -0.03]** | **-0.1\*\*\* (0.0); [-0.1, -0.03]** | **-0.1\*\*\* (0.0); [-0.1, -0.03]** |
| Sex | -0.01 (0.1); [-0.2, 0.2] | -0.02 (0.1); [-0.2, 0.2] | -0.03 (0.1); [-0.2, 0.2] |
| Education | -0.003 (0.0); [-0.01, 0.04] | -0.001 (0.0); [-0.04, 0.04] | 0.001 (0.0); [-0.04, .04] |
| Practice effect | 0.04 (0.1); [-0.1, 0.1] | 0.04 (0.1); [-0.1, 0.1] | 0.1 (0.1); [-0.1, 0.2] |
| Medication use at some visits | -0.1 (0.1); [-0.4, 0.1] | --- | 0.03 (0.1); [-0.2, 0.3] |
| Medication use at all visits | -0.1 (0.2); [-0.4, 0.3] | --- | -0.01 (0.1); [-0.3, 0.3] |
| Risk factor group x Amyloid group | 0.2 (0.2); [-0.2, 0.7] | -0.2 (0.2); [-0.7, 0.2] | -0.3 (0.3); [-0.9, 0.3] |
| Centered visit age x Risk factor group | -0.004 (0.0); [-0.02, 0.02] | -0.01 (0.0); [-0.03, 0.01] | -0.01 (0.0); [-0.04, .02] |
| Centered visit age x Amyloid group | -0.01 (0.0); [-0.03, 0.01] | -0.01 (0.0); [-0.03, 0.01] | -0.01 (0.0); [-0.03, .01] |
| Centered visit age x Amyloid group x Risk factor group | -0.01 (0.0); [-0.04, 0.03] | -0.01 (0.0); [-0.04, 0.03] | -0.02 (0.0); [-0.1, 0.03] |

\*\*\* $p \leq .001$ ; \*\* $p \leq .01$ ; \* $p < .05$ ; p-values for fixed effect coefficients were calculated using asymptomatic properties of the estimates

**Figure 2 Legend.**
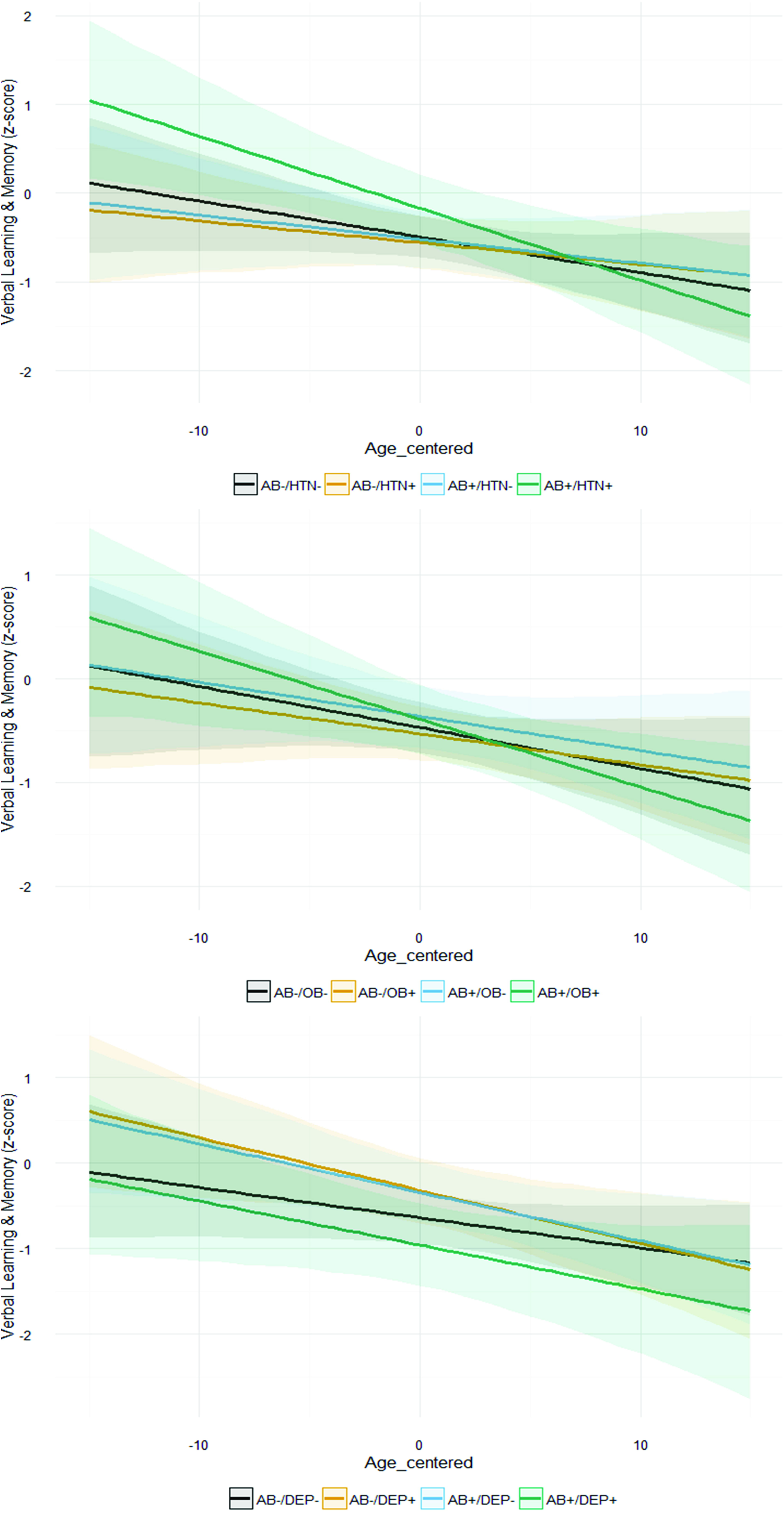
Graphs depict Verbal Learning & Memory z-scores on the y-axis, age at each visit (centered on mean age) on the x-axis, and estimated slopes for four beta-amyloid/risk factor groups adjusted for covariates of age at biomarker visit, sex, education, and practice effects. Risk factor groups are determined based on status across all visits available (negative = risk factors absent at all visits; positive = risk factor present at one or more visits). The top figure depicts the estimated slope for beta-amyloid negative and hypertension negative (black; *n* =93), beta-amyloid negative and hypertension positive (orange; *n* =52), beta-amyloid positive and hypertension negative (blue; *n* =37), and beta-amyloid positive and hypertension positive (green; *n* =25). The middle figure depicts the estimated slope for beta-amyloid negative and obesity negative (black; *n* =62), beta-amyloid negative and obesity positive (orange; *n* =83), beta-amyloid positive and obesity negative (blue; *n* =33), and beta-amyloid positive and obesity positive (green; *n* =29). The lower figure depicts the estimated slope for beta-amyloid negative and depression negative (black; *n* =123), beta-amyloid negative and depression positive (orange; *n* =22), beta-amyloid positive and depression negative (blue; *n* =50), and beta-amyloid positive and depression positive (green; *n* =12).

Additional analyses were conducted to examine the effect of antihypertensive medication use on these results. Nineteen percent (*n* =40) were treated with an antihypertensive medication. Of the sample of *n* =207, *n* =143 were non-hypertensive/non-treated, *n* =29 were non-hypertensive/treated, *n* =24 were hypertensive/non-treated, and *n* =11 were hypertensive/treated. The overall three-way interaction among hypertension/treatment status x amyloid status x visit age did not account for a statistically significant amount of variance in Verbal Learning & Memory performance (*χ*^*2*^(*3*) = 4.86, *p* = .18; see Supplemental Table 1). Although the omnibus interaction term was non-significant, evaluation of the model coefficients suggests that the hypertensive/untreated group with amyloid positivity exhibited greater age-related memory decline compared to the reference group (non-hypertensive/non-treated) *β* = −0.08, *t* = −2.11, *p* = .03), whereas the other groups did not differ from the reference group (see Supplemental Table 1).

#### Obesity

Likelihood ratio tests comparing the full and nested linear mixed-effects models indicated that the three-way interaction of obesity status x amyloid status x visit age was not statistically significant for the Verbal Learning & Memory factor (*χ*^*2*^*(1)* = 1.79, *p* = .18; Table 3); however, the direction of the effect was similar to that observed for hypertension (see Figure 1 [Middle]) and secondary analyses using cumulative obesity data (e.g., yes if obese at any visit and no if not obese at all visits) demonstrated a significant association between the three-way interaction of obesity status x amyloid status x visit age and Verbal Learning & Memory performance (*χ*^*2*^*(1)* = 3.89, *p* = .049; see Table 5). Statistical comparison of model coefficients indicated that those with amyloid positivity and obesity present at least one visit (Figure 2 [Middle green line]) exhibited greater decline than those with amyloid positivity without obesity at any visit (Figure 2 [Middle blue line]) (*β* = −0.03 (SE=.02), *p* = .08). Similar to analyses with hypertension, the three-way interaction of obesity status x amyloid status x visit age was not associated with performance on the Speed & Flexibility factor (*χ*^*2*^*(1)* = .90, *p* = .34; Table 4).

#### Depression

Likelihood ratio tests comparing the full and nested linear mixed-effects models indicated that the three-way interaction of depression status x amyloid status x visit age was not significantly associated with Verbal Learning & Memory (*χ*^*2*^*(1)* = 0.24, *p* = .62; see Table 3) or Speed & Flexibility (*χ*^*2*^*(1)* = 2.07, *p* = .15; see Table 4) performances. The two-way interaction between depression and amyloid status was significantly associated with Verbal Learning & Memory performance (*χ*^*2*^*(1)* = 4.55, *p* = .03; see Table 3), suggesting that verbal memory performance differed across one or more contrasts of the four groups defined by amyloid and depression status (see Figure 1 [Bottom]). The figure suggests worse performance in those with amyloid positivity and depression compared to other groups; however statistical comparison of model coefficients using the Wald test indicated that those with amyloid positivity and depression did not significantly differ from those with amyloid positivity without depression (*β* = −0.61 (SE=.38), *p* = .11), those with amyloid negativity with depression (*β* = −0.68 (SE=.38), *p* = .13), or those with amyloid negativity and no depression (*β* = −0.42 (SE=.38), *p* = .26). The two-way interaction between depression and amyloid status was not associated with Speed & Flexibility performance (*χ*^*2*^*(1)* = 0.38, *p* = .54; see Table 4).

Secondary analyses which replaced visit 2 depression status with cumulative depression status (e.g., yes if depressed at any visit and no if non-depressed at all visits) showed similar results in that the three-way interaction of depression status x amyloid status x visit age was not significantly associated with Verbal Learning & Memory performance (*χ*^*2*^*(1)* = 1.36, *p* = .24) or Speed & Flexibility performance (*χ*^*2*^*(1)* = .69, *p* = .40). Similarly, the two-way interaction between depression and amyloid status was significantly associated with Verbal Learning & Memory performance (*χ*^*2*^*(1)* = 10.45, *p* = .001; Table 5), but not with Speed & Flexibility performance (*χ*^*2*^*(1)* = 1.12, *p* = .29; Table 6).

Additional analyses were conducted to examine the effect of antidepressant medication use on these results. Twenty-eight percent (*n* =58) were treated with an antidepressant medication at visit 2. Of the sample of *n* =207, *n* =145 (70%) were non-symptomatic/non-treated, *n* =48 (23%) were non-symptomatic/treated, *n* =4 (2%) were symptomatic/non-treated, and *n* =10 (5%) were symptomatic/treated. Results of likelihood ratio tests comparing the full and nested linear mixed-effects models indicated that the three-way interaction among depression/treatment status x amyloid status x visit age did not account for a statistically significant amount of variance in Verbal Learning & Memory performance (*χ*^*2*^(*3*) = 4.67, *p* = .20; see Supplemental Table 1).

## DISCUSSION

In a sample of 207 late middle-aged adults enriched for Alzheimer’s disease risk, presence of hypertension or obesity moderated the relationship between beta-amyloid (on PET scan or in CSF) and longitudinal verbal memory, but not speed & flexibility, performance. These findings suggest that the presence of hypertension or obesity in midlife may exacerbate the subtle cognitive decline associated with beta-amyloid deposition. Presence of depression did not moderate the relationship between beta-amyloid and longitudinal cognitive performance. Although presence of hypertension and obesity moderated the relationship between amyloid and verbal memory performance, results from chi-square analyses indicated there were no differences in the proportion of participants classified as amyloid positive by hypertension, obesity, or depression status. These latter results suggest there may be an additive effect of amyloid pathology and the presence of these risk factors to accelerate cognitive decline. Further longitudinal follow-up is ongoing and needed to confirm if the individuals exhibiting greatest decline progressively worsen and develop clinical symptoms of dementia.

In this longitudinal study we operationalized risk factor status in two ways: presence of the risk factor at the earliest visit it was measured and presence of the risk factor at any visit. We observed similar patterns across both methods, but defining risk status based on presence at any visit produced results with generally stronger effects and clearer decline in the group with elevated amyloid and presence of hypertension or obesity. The stronger findings using the latter method may simply be due to the larger sample sizes of the risk factor groups (e.g., *n* = 77 were hypertensive at any visit vs *n* = 35 were hypertensive at visit 2), and therefore greater power to detect differences. These findings suggest that for future studies assessing effects of modifiable risk factors and Alzheimer’s disease biomarkers on cognition, examining presence of the risk factor at any visit (rather than at baseline or last visit only) may be useful.

Although we observed an interaction between amyloid and hypertension on cognitive trajectories, we did not find that hypertension was associated with elevated amyloid deposition. This is in contrast to some prior reports and may be because our sample was younger than some prior reports (e.g., mean age of ~60 in our study compared to mean ages of 69.4 (Nation *et al.*, 2013) and 86.9 (Hughes *et al.*, 2013)) or due to different analysis methods (e.g., Langbaum *et al.* (2012) demonstrated that systolic blood pressure and pulse pressure positively correlated with beta-amyloid distribution in frontal, temporal, and parietal regions, whereas we categorized participants in groups based on blood pressure and composite amyloid cutoffs). Furthermore, prior studies suggest that adults with hypertension who are not treated exhibit greater decline and are at increased risk of dementia when compared to those with hypertension who are treated (Gelber *et al.*, 2013; Gottesman *et al.*, 2014). Within our sample with hypertension, we did not observe a significant interaction among age at each visit x amyloid status x hypertension/treatment group; however, the sample sizes for these groups were small and this needs further evaluation in larger cohorts. Although the omnibus interaction term was nonsignificant, evaluation of the model coefficients suggests that the hypertensive/untreated group with amyloid positivity exhibited greater age-related memory decline than the non-hypertensive/non-medicated group, whereas the hypertensive/treated group did not differ from the non-hypertensive/non-medicated group.

Obesity has been less studied with regard to amyloid deposition and cognition compared to hypertension and depression. A couple of recent studies (Chuang *et al.*, 2016; Gottesman *et al.*, 2017) observed that higher body mass index in midlife was associated with greater amyloid deposition in late life, whereas our findings suggest that midlife obesity (as measured via waist circumference) is not associated with elevated amyloid burden measured during midlife. However, our study found that the presence of obesity at all study visits moderated the relationship between amyloid burden and cognitive decline, suggesting that the presence of both factors may accelerate cognitive decline. Additionally, in our sample there was overlap between the obese sample and the hypertensive sample, so it is possible that the relationship observed was driven by hypertension. More specific mechanisms associated with obesity, such as insulin resistance or diabetes, may need to be evaluated to parse out specific associations between obesity and amyloid burden.

Moreover, we observed that depressive symptoms did not moderate the relationship between beta-amyloid and cognitive decline. Many studies consistently observe that adults with depression tend to exhibit poorer cognitive performances than non-depressed adults, perhaps due to amotivation, fatigue, and concentration difficulties inherent in depression (Gotlib and Joormann, 2010). However, other studies suggest that older adults with depression are more likely to decline over time (Thomas and O’Brien, 2008). It has been debated as to whether this latter finding might be because depressive symptoms are a part of the prodromal symptoms of dementia. Our results suggest that depression does not exacerbate amyloid-related cognitive decline; further longitudinal follow-up as well as future studies on types of depressive symptoms endorsed (e.g., somatic, emotional, cognitive) and the onset period of symptoms (e.g., chronically depressed vs new onset depression in mid or late life) will help provide clarification on whether depression in midlife is a harbinger of clinical symptoms of dementia. Additionally, a smaller proportion of the sample endorsed clinically significant depressive symptoms compared to those who met criteria for hypertension or obesity; it is possible that participants with depression in the larger Wisconsin Registry for Alzheimer’s Prevention are less likely to participate in biomarker study procedures and therefore these results may reflect a smaller and biased sample of depressed individuals.

The current study adds to the literature by demonstrating that hypertension, and to a lesser extent obesity, moderates the relationship between amyloid and cognitive decline in middle-age. However, this study has several limitations that need to be considered when interpreting these results. First, these findings may not generalize to populations that are dissimilar to the current study cohort who are generally at higher risk for Alzheimer’s disease, well-educated, and Caucasian. Participants in this study have at most mild cognitive impairment and it is not certain that all participants who exhibit greater decline on neuropsychological measures will progress to a diagnosis of dementia; continued follow up of these participants is needed. Although these results are promising in that they suggest potential preventative strategies to mitigate effects of amyloid on cognition, future studies are needed to determine if treating these risk factors prevent or delay the onset of clinical symptoms of dementia.

## ACKNOWLEDGMENTS

We gratefully acknowledge the assistance of researchers and staff at the Wisconsin Registry for Alzheimer’s Prevention and Wisconsin Alzheimer’s Disease Research Center for assistance in recruitment and data collection. Most importantly, we thank the dedicated Wisconsin Registry for Alzheimer’s Prevention participants for their continued support and participation in this research.

